# Modeling driver cells in developing neuronal networks

**DOI:** 10.1101/260422

**Authors:** Stefano Luccioli, David Angulo-Garcia, Rosa Cossart, Arnaud Malvache, Laura Módol, Vitor Hugo Sousa, Paolo Bonifazi, Alessandro Torcini

## Abstract

Spontaneous emergence of synchronized population activity is a characteristic feature of developing brain circuits. Recent experiments in the developing neo-cortex showed the existence of *driver cells* able to impact the synchronization dynamics when single-handedly stimulated. We have developed a spiking network model capable to reproduce the experimental results, thus identifying two classes of driver cells: functional hubs and low functionally connected (LC) neurons. The functional hubs arranged in a clique orchestrated the synchronization build-up, while the LC drivers were lately or not at all recruited in the synchronization process. Notwithstanding, they were able to alter the network state when stimulated by modifying the temporal activation of the functional clique or even its composition. LC drivers can lead either to higher population synchrony or even to the arrest of population dynamics, upon stimulation. Noticeably, some LC driver can display both effects depending on the received stimulus. We show that in the model the presence of inhibitory neurons together with the assumption that younger cells are more excitable and less connected is crucial for the emergence of LC drivers. These results provide a further understanding of the structural-functional mechanisms underlying synchronized firings in developing circuits possibly related to the coordinated activity of cell assemblies in the adult brain.

**Author Summary:** There is timely interest on the impact of peculiar neurons (*driver cells*) and of small neuronal sub-networks (*cliques*) on operational brain dynamics. We first provide experimental data concerning the effect of stimulated driver cells on the bursting activity observable in the developing entorhinal cortex. Secondly, we develop a network model able to fully reproduce the experimental observations. Analogously to the experiments two types of driver cells can be identified: functional hubs and low functionally connected (LC) drivers. We explain the role of hub neurons, arranged in a clique, for the orchestration of the bursting activity in control conditions. Furthermore, we report a new mechanism, which can explain why and how LC drivers emerge in the structural-functional organization of the enthorinal cortex.

## Introduction

Coordinated neuronal activity is critical for a proper development and later supports sensory processing, learning and cognition in the mature brain. Coordinated activity represents also an important biomarker of pathological brain states such as epilepsy [1]. It is therefore essential to understand the circuit mechanisms by which neuronal activity becomes coordinated at a population level. A series of experimental results indicates that non-random features are clearly expressed in cortical networks [2–4], in particular neuronal sub-networks, termed *cliques*, have been shown to play a fundamental role for the network activity and coding both in experiments [5–9] as well as in models [10–13].

The identification of these small highly active assemblies in the hippocampus [5] and in the cortex [6–8] poses the question if these small neuronal groups or even single neurons can indeed control the neural activity at a mesoscopic level. Interestingly, it has been shown that the stimulation of single neurons can affect population activity in vitro as well as in vivo [14–23]. The direct impact of single-neurons on network and behavioral outputs demonstrates the importance of the specific structural and functional organization of the underlying circuitry. Neurons having such a network impact were recently termed *operational hubs* [24] or *driver cells* [23]. It is thus critical to understand how specific network structures can empower single driver cells to govern network dynamics. This issue has been addressed experimentally in some cases. More specifically, in the developing CA3 region of the hippocampus, single GABAergic hub neurons with an early birthdate were shown to coordinate neuronal activity. These cells have a high functional connectivity degree, reflecting mainly the fact that they are activated at the onset of Giant Depolarizing Potentials (GDPs), as well a high effective connectivity degree [16]. This therefore represents a simple case where the circuit mechanism, promoting a cell to the role of hub, is due to their exceptional number of anatomical links. But the picture can be quite different in other brain regions, as recently demonstrated in the developing Entorhinal Cortex (EC) [23], where the driver cell population comprises both cells with a high functional out-degree, as well as low functionally connected (LC) cells.

In order to understand the circuit mechanisms by which even a LC cell can influence population bursts we have upgraded and modified a network model based on excitatory leaky integrate-and-fire (LIF) neurons [11], previously developed to reproduce the functional properties of hub neurons in the developing hippocampal circuit [16]. In such a model the *population bursts* (PBs), corresponding to GDPs in neonatal hippocampus [25], were controlled by the sequential and coordinated activation of few functional hubs. Notably, the perturbation of one of these neurons strongly impacted the collective dynamics and brought even to the complete arrest of the bursting activity, similarly to what experimentally found for the developing hippocampus in [16]. The model described in this paper contains two main differences with respect to the hippocampal model [11]. Firstly, it comprises both inhibitory and excitatory neurons, to account for the fact that, even though GABA acts as an excitatory neurotransmitter at early postnatal stages, some more developed neurons have already made the switch to an inhibitory transmission at the end of the first postnatal week in mice (P8), where most experimental data was obtained [26–28]. Secondly, the developmental profile of the network is regulated only by the correlation between neuronal excitability and connectivity, while in [11] a further correlation was present.

This model nicely mimicked the experimental observations in the EC similarly displaying the presence of driver cells with both low and high functional connectivity. We will first compare a few examples of LC drivers impacting circuits’ dynamics both in the EC and in our model. Next, we will present a full characterization of the numerical model leading to a complete understanding of the mechanism underlying the PB generation and the impact of LC driver cells on population dynamics.

## Results

### Experimental Evidences of driver LC cells

The main experimental observation at the rationale of this work is the existence of *driver cells* (or Operational Hubs [24]) in the mice EC during developmental stage [23]. Driver cells have been identified using calcium imaging experiments and they were charaterized by the capability to impact network synchronization (namely, GDPs' occurence) when externally activated/stimulated through intra-cellular current injection. Two classes of driver cells were identified: (i) those with high directed functional connectivity out-degree, early activated and playing a critical role in the network synchronizations (*driver hub cells*) and (ii) those recruited only in the later stages of the synchronization build up, which therefore are low functionally connected (*driver LC cells*).

The experimental setup used to identify, target and probe the single-handedly impact of neurons on spontaneous EC synchronization is schematized in Fig. 1 (a.E) and Fig. S1. In brief, the functional connectivity of the cells has been measured during the spontaneous activity session, which preceded the single neurons’ stimulation session, both lasting two minutes. A directed functional connection from neuron A to B was established whenever the firing activity of A significantly preceded the one of neuron B (more details can be found in *Methods*). The functional out-degree 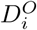 of a neuron *i* corresponded to the percentage of imaged neurons which were reliably activated after its firing. Neurons in the 90% percentile of the connectivity distribution were classified as hub neurons early activated in the network synchronization.

**Figure 1.**
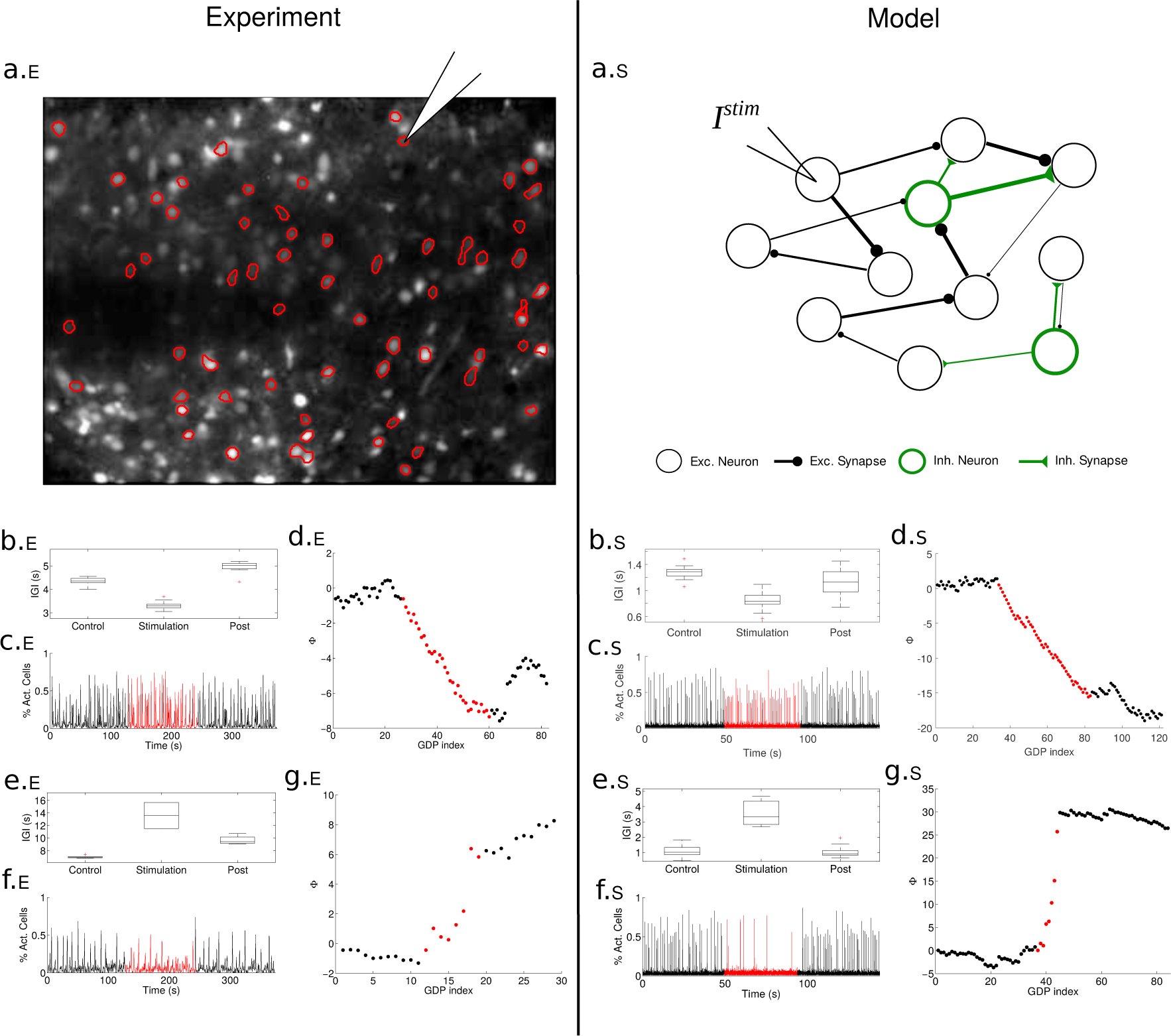
Impact of single-handedly stimulation of LC drivers on the collective dynamics of the Enthorinal Cortex and of the neuronal network model. The left column (x.E) refers to experimental measurements taken in slices of EC, while the right column (x.S) to the numerical results. Experiment. The first panel (a.E) presents the image of a slice loaded with the calcium sensor where the active neurons are encircled and the pipette indicates the neuron targeted for intracellular stimulation. Data for a neuron with functional out-degree *D*^*O*^ 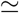 7% are reported in (b-d.E), data for a neuron (from a different slice) with *D*^*O*^ 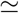 8% are shown in (e-g.E). Panels (b.E) and (e.E) are boxplot of the IGIs for each experimental phase. Panels (c.E) and (f.E) represent the fraction of recruited cells participating in the GDP. During the stimulation period (red curves) a single cell is stimulated with a frequency *v*_*S*_ = 0.33 Hz (*v*_*S*_ = 0.14 Hz) for the first (second) neuron according to the protocol discussed in the text. Panels (d.E) and (g.E) report the phase Φ of the GDP as a function of time (ticked by the GDP index), specifically the difference between the number of expected versus observed GDP based on the pre-stimulation frequency (see *Methods*). The average IGI was 4.2 s (7.8 s) in the pre-stimulation phase, becoming 3.2 s (14 s) during the stimulation period for the first (second) neuron. **Model**. The first row (a.S) displays a cartoon of the performed single neuron stimulation experiment in the network model, where inhibitory (*inh*.) and excitatory (*exc*.) neurons and synapses are marked in black and green respectively. Panels (b-d.S) refer to the excitatory driver LC cell *el*_1_ which was silent in control condition and once stimulated with a current *I*^stim^ = 15.135 mV was able to enhance population dynamics. Panels (e-g.S) refer to the excitatory driver LC cell el_2_ which was active in control condition with D^O^ = 3% and once stimulated with a current *I*^stim^ = 15.435 mV depressed the PB activity.

The protocol used for probing the impact of single neurons on the network dynamics was organized in three phases, each of two minutes duration: (1) a pre-stimulation resting period; (2) a stimulation period, during which a series of supra-threshold current pulses at a specific frequency *v*_*S*_ (of the order of the GDPs frequency) have been injected into the cell; (3) a final recovery period, where the cell is no more stimulated. The impact on the network activity of the single-handed stimulation was measured by employing two indicators (see *Methods* for more details) : (i) the variation of the average Inter-GDP-Interval (IGI) during the stimulation phase with respect to the resting period; (ii) the shift of the IGI phase Φ, as defined in *Methods* Eq. (1) and in [29], induced by the stimulation with respect to the prestimulation period. At a population level the stimulation could have an inhibitory (excitatory) effect corresponding to a slow-down (acceleration) of the GDP frequency associated with an increase (decrease) of the measured IGI and with a positive (negative) phase shift.

Two examples of driver LC cells, with 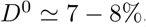, are reported in Fig 1 in the panels (b-d.E) and (e-g.E). In the first case, upon stimulation the network dynamics accelerated, as testified by the decrease of the average IGI (Fig 1 (b.E)) and by the negative instantaneous phase shift of GDPs (Fig 1 d.E). In the second case, the stimulation led to a pronounced slow down of the average network activity (as shown in Fig. 1 (e.E)) together with an increase of the instantaneous phase with respect to control conditions (Fig. 1 (g.E)). In both cases the removal of the stimulation led to a recovery of the dynamics similar to the control ones.

A further extreme case of a silent cell, i.e. not spontaneously active and therefore with a zero (out-degree) functional connectivity, is shown in Fig.2. This cell, when stimulated with different stimulation frequencies *v*_*S*_, revealed opposite effects on the network behaviour. At lower stimulation frequency (*v*_*S*_ = 0.33 Hz) the cell activity induced an acceleration of the population dynamics (see Fig. 2 (a-c.E)), while at higher stimulation frequency (*v*_*S*_ = 1 Hz) of the same neuron we observed a slowing down of the network dynamics (see Fig. 2 (d-f.E)).

**Figure 2.**
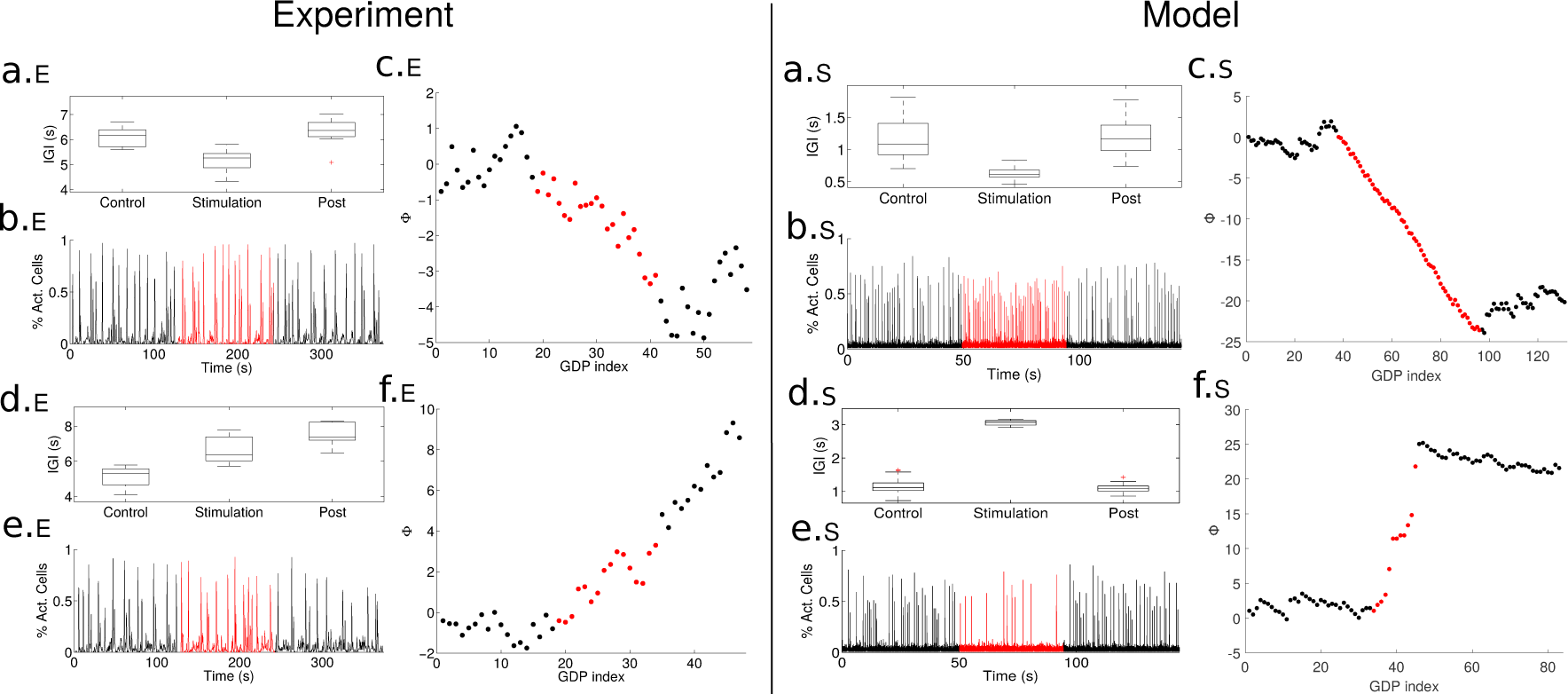
The same driver LC cell upon different stimulations can enhance or depress the population activity. The left column (x.E) refers to experimental measurements taken in slices of enthorinal cortex, while the right column (x.S) to the numerical results. Panels are similar to Fig. 1 but reporting the case in which the network acceleration (a-c.E and a-c.S) and slowing down (d-f.E and d-f.S) are observable by stimulating the same neuron at different frequencies. The experiment and the simulation refer to neurons with zero functional out-degree. **Experiment**. Panels (a-c.E) ((d-f.E)) are obtained with *v*_*S*_ = 0.33 Hz (*v*_*S*_ = 1.0 Hz). Namely, the average IGI varied from 6.08 s in control conditions to 5.14 s during stimulation with *v*_*S*_ = 0.33 Hz, and from 5.36 s to 6.68 s with *v*_*S*_ = 1.0 Hz. **Model**. All the panels refer to the driver LC cell el_3_ connected in output to the eli neuron discussed in panels (b-d.E) in Fig. 1. The results refer to the stimulation of the same neuron with two different currents, namely panels (a-c.S) refer to *I*^stim^ = 15.42 mV and (d-f.S) to current *I*^stim^ = 15.9 mV.

### Numerical Evidences of driver LC cells

In order to mimic the impact of single neurons on the collective dynamics of a neural circuit, we considered a directed random network made of N LIF neurons [30,31] composed of excitatory and inhibitory cells and with synapses regulated by short-term synaptic *depression* and *facilitation*, analogously to the model introduced by Tsodyks-Uziel-Markram (TUM) [32] (see *Methods* for more details). As shown in [32–34], these networks exhibit a dynamical behavior characterized by an alternance of short periods of quasi-synchronous firing (PBs) and long time intervals of asynchronous firing, thus resembling cortical GDPs’occurrence in early stage networks. Similarly to the modeling reported in [11], we considered neuronal intrinsic excitabilities negatively correlated with the total connectivity (in-degree plus out-degree) (for more details see *Definition of the Model* in *Methods* and Fig. S2). The introduction of these correlations was performed to mimic developing networks, where both mature and young neurons are present at the same time associated to a variability of the structural connectivities and of the intrinsic excitabilities. Experimental evidences point out that younger cells have a more pronounced excitability, most likely due to the fact that their GABAergic inputs are still excitatory [35–37], while mature cells exhibit a higher number of synaptic inputs and they do receive inhibitory or shunting GABAergic inputs [16,38]. The presence of inhibition and facilitation are the major differences from the model developed in [11] to simulate the dynamics of hippocampal circuits in the early stage of development, justified by the possible presence of mature GABAergic cells in the network.

Using this network model, we studied the effect of single neuron current injection /^stlm^ on network dynamics, thus altering the average firing frequency of the neuron during the stimulation time, similarly to what done in the experiments. In the numerical investigations, at variance with the experiments, the stimulation delivered to the neurons is an unique supra-threshold step of duration of 50 seconds. In Fig. 1 two representative driver LC cells are reported for comparison with the experiments. The first cell (panels (b-d.S) of Fig. 1) was a silent neuron in control conditions (therefore with *D*^*O*^ = 0), that once stimulated could enhance of 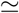 30 % the PB emission, thus leading to a decrease of the instantaneous phase Φ with respect to control condition. Panels (e-g.S) refer to a second neuron characterized by a low functional output connectivity, namely *D*^*O*^ = 3%, whose stimulation led to a depression in the PB frequency (as shown in panels (e.S) and (f.S)) joined to an increase of the instantaneous phase of the network events with respect to control conditions (as shown in panel (g.S)). These results compare quite well with the experimental findings reported in the same figure.

Furthermore, analogously to what found in the experiment, Fig. 2 (a-f.S) shows a silent neuron in control condition that once stimulated could lead to both enhancement or depression of the population activity depending on the level of injected current during stimulation.

A full characterization of the network model concerning the impact on the network dynamics of each single neuron stimulation in relation to neuronal type, current injected and functional connectivity is detailed below.

### Impact of single neuron stimulation and deletion on network dynamics

In order to explore the full dynamical range associated to the impact of single neuron stimulation on the network dynamics, we examined the response of the model network to two types of single neuron perturbations, i.e. *single neuron deletion* (SND) and *single neuron stimulation* (SNS) by employing the protocols introduced in [11]. In particular, the SND experiment consisted in recording the activity of the network in a fixed time interval *Δt* = 84 s when the considered neuron was removed from the network itself. While, the stimulation of the single neuron (SNS) was performed with a step of DC current of amplitude *I*^stim^ for a time window *Δt* = 84 s. The recording of the activity in control condition was lasting 84 s as well, in order to compare directly the number of observed PBs during control and perturbation period. In particular, we tested the response of the network to a broad range of stimulation amplitudes varying from 14.5 mV (slightly below the firing threshold for an isolated neuron *V*_*th*_ = 15 mV, see *Methods*) to 18.0 mV with a step of 0.015 mV, inducing in the stimulated neuron a maximal firing frequency of 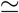 70 Hz. Typically the stimulated neuron fired with a frequency much higher than the frequency of neurons under control conditions (i.e. in absence of any perturbation). As an example, for a stimulation current *I*^stim^ = 15.90 mV the targeted neuron fired at a frequency *ν* 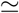 32 — 36 Hz well above the average (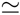 3 Hz) and the maximal (22 Hz) frequency of all neurons in control conditions. The SND represented an extreme version of the SNS, where the neuronal removal corresponded to the injection of an hyperpolarizing current inhibiting the neurons from firing spontaneously or in response to any synaptic input.

In both SNS and SND experiments, the impact of single neuron perturbation on the collective dynamics, was measured by the variation of the PB frequency relative to control conditions. In general, we have classified a neuron as a driver cell whenever upon stimulation it is able to modify the PB frequency at least or more than 50% with respect to control conditions. In the specific, in analogy to what done in [23], for SNS experiments we considered both enhancement and decrease in the PB activity. On the other hand, SND allowed us to directly identify the driver neurons which are fundamental for the PB build-up. Therefore in this case we limited to consider those cells, whose SND led to a population burst decrease at least or larger than 50%.

Fig. 3 (a-b) reports a comparison of the impact of SND and SNS (with representative injected current of 15.90 mV) on the PB activity. The removal of any of the four neurons labeled as *ih_1_,eh_1_,eh_2_,eh_3_* was able to arrest completely the bursting dynamics within the considered time window, while in other two cases (for neurons *ih*_2_ and *eh*_4_) the activity was reduced of 60% with respect to the one in control conditions. For clarity, the used labels *i/e* stand for inhibitory/excitatory and *h* for hub, as we will show later this is related to the functional role played by these cells. For all the other neurons, the SND manipulation induced a non relevant modification in the number of emitted PBs, within the variability of the bursting activity in control conditions (Fig. 3 (a)).

**Figure 3.**
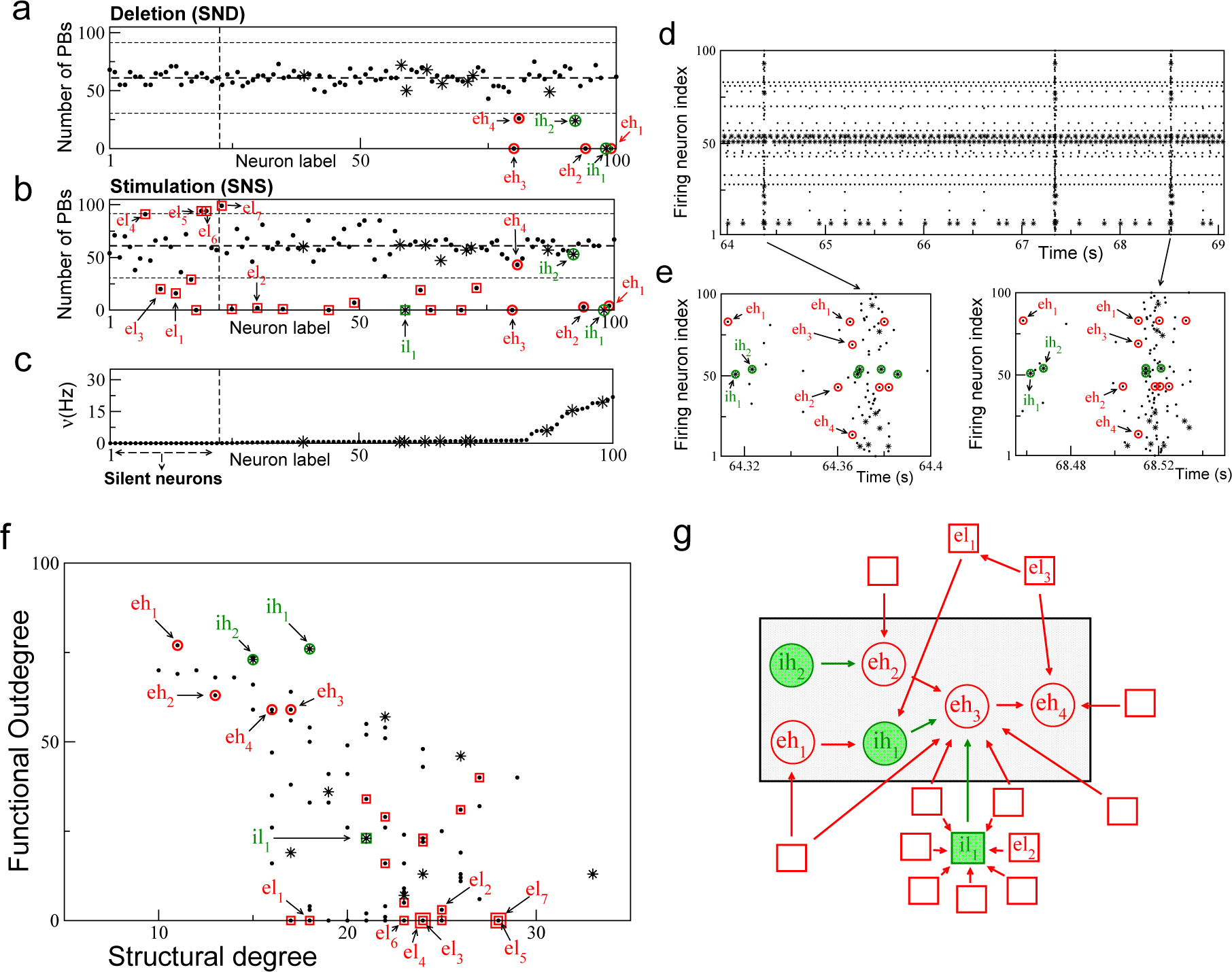
Model - Network response to single neuron stimulation. (a-b) Response of the network to SND and SNS, respectively. These panels report the number of PBs, recorded during SND (SNS) experiments versus the labels of the removed (stimulated) neuron, ordered accordingly to their average firing rates *ν* under control conditions (shown in panel (c)). Inhibitory neurons are marked with asterisks. In this representative SNS experiment each neuron was stimulated with a DC step *I*^stim^ = 15.90mV for a time interval Δ*t* = 84 s. The central horizontal dashed line shows the average number of PBs emitted in control conditions within an interval Δ*t* = 84 s, while the lower and upper horizontal dashed lines mark the 50% variation. The vertical dashed line separates firing neurons (on the right side) from silent neurons (on the left side) in control conditions. (d) Raster plot of the network activity. (e) Close ups of population bursts representative of the two principal routes (i.e the firing sequence of hub cells): an instance of the main route is reported in the left panel, while the second most frequent route is displayed in the right panel. Note that neurons are labeled accordingly to their index and are not ordered as in panels (a-c). (f) Scatter plot showing the functional out degree *D*^*O*^ of the neurons versus their total structural degree *K*^*T*^. A double square marks two neurons with overlapping properties. (g) Sketch of the structural connections among driver hub neurons and LC1 driver cells, the gray shaded rectangle highlights the clique of the hub cells. In all the panels, circles (squares) denote hub (LC) driver cells, while the green (red) symbols refer to inhibitory (excitatory) driver cells.

The SNS confirmed that the neurons *ih*_1_, *eh*_1_, *eh*_2_, *eh*_3_ were capable to arrest the collective dynamics. Neurons *eh*_4_ and *ih*_2_ poorly impacted PB dynamics for the reported injected current, although for different values of *I*^stim^ they were able to strongly influence the network dynamics (as shown in the subsection *Tuning of PBs frequency upon hubs’ and driver LC cells’ stimulations).* At variance from what found in a purely excitatory network [11], the SNS revealed also the presence of other 18 driver cells not identified by the SND capable to impact the occurence of PBs in the network (Fig. 3 (b)).

For an equivalent random network, without any imposed correlation, SNS or SND affected the dy-namics in a neglibile way producing a maximal variation of the bursting activity of 25-30 % with respect to the control conditions (see Fig. S3 (a-b)).

To summarize, the presence of correlations among the neuronal intrinsic excitabilities and the corresponding structural connectivities was crucial to render the network sensible to single neuron manipulation. Differently from purely excitatory networks where SNS and SND experiments gave similar results, the inclusion of inhibitory neurons in the network promoted a larger portion of neurons to the role of drivers, and their properties will be investigated in the following.

### Connectivity and excitability of the driver cells

The role played by the neurons in the simulated network was elucidated by performing a directed functional connectivity (FC) analysis. In the case of the spiking network model, in order to focus on the dynamics underlying the PB build-up, the FC analysis was based on the first spike fired by each neuron in correspondence of the PBs. An equivalent information was provided in the analysis of the EC by considering the calcium signal onset to calculate the directed functional connectivity. The six neurons playing a key role in the generation of the PBs (*eh*_1-4_, *ih*_1-2_), were characterized by high values of functional out-degree, namely with an average functional degree *D*^*O*^ = 68%±8%, ranking them among the 16 neurons with the highest functional degree. Given the high functional out-degree and their fundamental role in the generation of the PBs (as shown by the SND in Fig. 3 (a)), we identified these neurons as *driver hub cells*. The high value of *D*^*O*^ reflected their early activation in the PB, thus preceding the activation of the majority of the other neurons.

Next, we examined the structural degree of the neurons, specifically we considered the total structural degree *K*^*T*^, which is the sum of the in-degree and out-degree of the considered cell. As shown in Fig. 3 (f), we observed an anti-correlation among *D*^*O*^ and *K*^*T*^ where neurons with high functional connectivity are typically less structurally connected than LC neurons. This was particularly true for the six driver hubs, previously examined, since they were characterized by an average *K*^*T*^ = 15 ± 3, well below the average structural connectivity of the neurons in the network (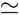 20).

Concerning the excitability, the six driver hubs despite being in proximity of the firing threshold (slightly above or below) as shown in Fig. S4 (a), they were among the 25% fastest spiking neurons in control condition, (as shown in Fig. 3 (c)). In particular, the three neurons *eh*_1_, *eh*_2_, *ih*_2_ were supra-threshold, while neurons *eh*_3_, *eh*_4_, *ih*_1_ were slightly below the threshold. When embedded in the network their firing activity was modified, in particular three couples of neurons with similar firing rates can be identified, namely (*eh*_1_,*ih*_1_), (*ih*_2_,*eh*_2_) and (*eh*_3_,*eh*_4_), as reported in Table I. The direct structural connections present among these couples (see also Fig. 3 (g)) could explain the observed firing entrainments, as discussed in details in the next subsection. When compared to the other hub neurons, the much lower activity of (*eh*_3_, *eh*_4_), corresponding to twice the average frequency of the PBs in control condition, was related to the fact that these two neurons fired only in correspondence of the ignition of collective events like PBs and *aborted bursts* (ABs), the latter being associated to an enhancement of the network activity but well below the threshold we fixed to detect PBs. This will become evident from the discussion reported in the subsection *Synaptic resources and population bursts*.

**Table 1.** Properties of driver hub cells in control condition. For each driver hub cell (*ih*_1_, \**ih*_2_, *eh*_1_, *eh*_2_, *eh*_3_, *eh*_4_) the columns report the intrinsic excitability (*I*^b^), the average spiking frequency in control conditions (*v*), the structural out-degree (*K*^O^) and in-degree (*K*^I^), the functional out-degree (*D*^O^) and in-degree (*D*^I^).

| Driver hub cell | $I^b$ (mV) | $\nu$ (Hz) | $K^O$ | $K^I$ | $D^O$ | $D^I$ |
| --- | --- | --- | --- | --- | --- | --- |
| $ih_1$ | 14.91 | 19.5 | 11 | 7 | 76% | 1% |
| $ih_2$ | 15.02 | 15.5 | 6 | 9 | 73% | 3% |
| $eh_1$ | 15.32 | 20.5 | 6 | 5 | 77% | 0% |
| $eh_2$ | 15.23 | 16.3 | 7 | 6 | 63% | 11% |
| $eh_3$ | 14.93 | 1.3 | 7 | 10 | 59% | 12% |
| $eh_4$ | 14.99 | 1.3 | 9 | 7 | 59% | 12% |

As already mentioned, besides the six driver hubs, the SNS experiments revealed the existence of a different set of 18 drivers, whose activation also impacted the population dynamics, although they had no influence when removed from the network and therefore they were not relevant for the PBs build up. These neurons represented in Fig. 3 with squares were characterized by a low FC, namely *D*^*0*^ = 13%±15%. Therefore, we have termed them *driver LC cells* representing the ones which reproduced the behaviour of the driver LC cells identified in the EC (see Fig. 1 and 2 and reference [23]). In the following we will refer to them as *el*… or *il*_1_ according to the fact that they are excitatory or inhibitory neurons, respectively (note that only one LC driver was inhibitory). As shown in Fig. 3 (c), LC drivers were not particularly active (with firing frequencies below 1 Hz in control conditions) and in some cases they were even silent. Notably, under current stimulation they could in several cases arrest PBs or strongly reduce/increase the activity with respect to control conditions as shown in Fig. 3 (b) for a specific level of current injection and also as discussed in detail in the following sections.

Compared to the drivers hubs, driver LC cells had a lower degree of excitability (essentially they were all sub-threshold, see Fig. S4 (a)), which resulted in a later recruitment in the synchronization build up, and as a consequence in a lower functional out-degree. Therefore, driver LC cells were not necessary for the generation of the PBs, playing the role of followers in the spontaneous network synchronizations. As shown in Fig. 3 (f), driver LC neurons were charaterized by a higher structural connectivity degree *K*^*T*^ with respect to driver hubs, namely *K*^*T*^ = 23 ± 3, and the most part of them were structurally targeting the hub drivers either directly (i.e. path length one) or via a LC driver (i.e. path length two, centered on a LC driver). In Fig. 3 (f), the two groups of drivers, hubs and LC cells, can be easily identified as two disjoint groups in the plane (*K*^*T*^, *D*^0^). These results indicated that driver hubs are not structural hubs, while the low functional connectivity neurons are promoted to their role of drivers due to their structural connections. This latter aspect will be exhaustively addressed in subsection *Tuning of PBs frequency upon hubs’ and LC cells’s stimulation*.

### Statistical analysis

The reported results are statistically significant, as we have verified by analyzing fifteen different realizations of the network. In particular, we used the same distributions for the intrinsic excitabilities, synaptic parameters and structural connectivities. The parameter values were taken from random distributions with the same averages and standard deviations as defined in *Definition of the model* in *Methods*. Furthermore, in all the numerical experiments we kept fixed the size of the network (*N* = 100), the number of excitatory/inhibitory neurons (*N*_*e*_ = 90, *N*_*i*_ = 10), the average in-degree, and all the other constraints specified in *Definition of the model* in *Methods*. In six networks we found no bursting dynamics or number of bursts too small to be significant. While, in the remaining nine network PBs were always present and we could perform significant SND/SNS experiments on all the neurons in each network. This analysis allowed us to identify driver hub cells and driver LC cells in all these networks, with characteristic similar to the ones found in the network analysed in detail in the paper. In particular, we have identified for each network a number of hub cells ranging from two to eight with an average value 5 ± 2, and a number of LC cells ranging from 1 to 27 with an average value 12 ± 9 (apart a peculiar single network where we found just one hub and one LC cell). By examining the nine networks displaying bursting dynamics we found the presence of inhibitory cells among the hubs in three cases and among the LC driver cells in six cases (with numbers ranging from one to four). As general features, we observed that hub driver cells were characterized by a high intrinsic excitability and a low structural connectivity: namely, *I*^b^ was in the range [14.55 : 15.42] mV (with average 15.0 ± 0.2 mV and 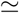 37% of the hubs supra-threshold), while the total connectivity *K*^*T*^ was in the range [6 : 31] (with average 16 ± 3 and a single hub with *K*^*T*^ = 31). On the other hand, the LC drivers were characterized by a low intrinsic excitability in the range [14.55 : 15.03] mV (with average 14.7 ± 0.1 mV and 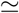 99% of the LC cells below the firing threshold) and by a high *K*^*T*^ in the range [14 : 32] (with average 22 ± 4 and a single LC driver cell with *K*^*T*^ =14).

### Functional clique of excitatory and inhibitory neurons

In order to deepen the temporal relationship among neural firings leading to a PB, we examined the spikes emitted in a time window of 70 ms preceding the peak of synchronous activation (see *Methods* for details). The cross correlations between the timing of the first spike emitted by each hub driver neuron during the PB build up are shown in Fig. S5 (Upper Sequence of Panels). The cross correlation analysis demonstrated that the sequence of activation of the neurons was *eh*_1_ → *ih*_1_ → *ih*_2_ → *eh*_2_ → *eh*_3_ → *eh*_4_. The labeling previously assigned to these neurons reflected such an order. A common characteristic of these cells was that they had a really low functional in-degree *D*^*I*^ as reported in Table I indicating that they are among the first to fire during the PB build-up. In particular, *eh*_1_ had a functional in-degree *D*^*I*^ zero, revealing that it was indeed the firing of this neuron to initialize all the bursts and therefore it could be considered as the *leader* of the clique.

A detailed inspection of the firing times, going beyond the first spike event, revealed the existence of more than one firing sequence leading to the collective neuronal activation: i.e. the existence of different routes to PBs. This is at variance with what found in [11] for a purely excitatory network, where only one route was present and all the PBs were preceded by the same ordered sequential activation of the most critical neurons. In particular, the neuron *eh*_1_ fired twice before the PBs (see Fig. 3 (e)), usually in-between the firing of *eh*_2_ and that of the pair (*eh*_3_, *eh*_4_), and this represents the main route, occurring for 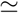 85% of the PBs. Along the second route (present only for the 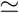 7% of the PBs), *eh*_1_ was firing the second time at the end of the sequence. The neuron *eh*_1_ fired essentially by following its natural period 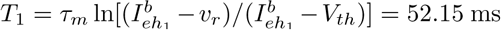, and its second occurrence in the firing sequence depended on the delay among the firing of the other neurons. As a matter of fact we verified that the elimination of the second spike emitted by eh_1_ from the network dynamics didn’t prevent, and didn’t delay, the onset of the PB and had only a marginal effect on the firing of a very limited number of neurons in the PB. Therefore we can conclude that it is not essential to the PB build up. The two routes leading to the PB build-up are shown in Fig. 3 (e).

To observe a PB the six driver hubs should fire not only in an ordered sequence, as shown in Fig 3 (e), but also with defined time delays, their average values with the associated standard deviations are reported in Table S1 for the two principal routes. These results clearly indicate that the six driver hubs are arranged in a *functional clique* whose activation was crucial for the PB build-up. In the period between the occurence of two PBs, the driver hubs in the clique could be active, but in that case they did not show the precise sequential activation associated to the main and secondary route, see the out-of-burst results reported in the Lower Sequence of Panels in Fig. S5. A remarkable exception is represented by the case of the ABs, in that case PBs are not triggered despite the presence of the right temporal activation of all the hubs in the clique, due to the lack of synaptic resources (as discussed in details in subsection *Synaptic resources for population bursts*). Out of PBs and ABs, we registered clear time-lagged correlations only for those neuronal pairs sharing direct structural connections (shown in Fig. 3 (g)): namely, e*h*_1_ → *ih*_1_, *ih*_2_ → *eh*_2_ and *eh*_2_ → (*eh*_3_, *eh*_4_). The firing delays of these neuronal pairs were not particularly altered also out of burst with respect to those measured during the burst build-up and reported in Table S1.

As shown in Fig. 3 (g), the *eh*_3_ neuron represented the cornerstone of the clique, receiving the inhibitory input coming from the structural pair (*eh*_1_, *ih*_1_) and the excitatory one from the pair (*ih*_2_, *eh*_2_), with the activity of the neurons within each pair perfectly frequency locked. More specifically, *eh*_1_ entrained the activity of *ih*_1_ (below threshold in isolation) so that both neurons before a PB fired with a period quite similar to the natural period of *eh*_1_. The other pair (*ih*_2_, *eh*_2_) was controlled by the inhibitory action of *ih*_2_ that slowed down the activity of *eh*_2_, whose natural period was 60.6 ms, while before a PB *ih*_2_ and *eh*_2_ both fired with a slower period, namely 72 ± 2 ms.

As it will be explained in details in the next two subsections, the two requirements to be fulfilled for the emergence of PBs are the availability of sufficient synaptic resources at neurons eh_3_ and *eh*_4_ and the coordinated activation of *eh*_1_ (and *ih*_1_) with the pair (*ih*_2_, *eh*_2_), in the absence of any synaptic connection between the two pairs.

### Synaptic resources for population bursts

Next we analyzed the relation between the evolution of synaptic resources in the driver hub cells and the onset of the PB. The availability of synaptic resources was measured by the effective efferent synaptic strength *X*^*OUT*^ as defined in Eq. (8). In particular, we will consider the available resources only for the hub neurons *eh*_3_ and *eh*_4_ which were the last neurons of the clique to fire before the PB ignition. We have examined only these two hub neurons, because whenever *eh*_3_ and *eh*_4_ fired, a burst or an AB was always delivered.

Neurons *eh*_3_, *eh*_4_ were receiving high frequency excitatory inputs from *eh*_2_ (although the natural firing of *eh*_2_ was slowed down by the incoming inhibition of *ih*_2_) and high frequency inhibitory inputs from *ih*_1_ (entrained by the *eh*_1_, the neurons with highest firing frequency in the network). This competitive synaptyc inputs resulted in a rare activation of *eh*_3_ compared to the higher frequency of excitatory inputs arriving from *eh*_2_. The period of occurrence of the ABs was comparable to the average interval between PBs (namely, *T*_*pb*_ = 1.4 ± 1.0 s) and ABs were preceded by the sequential activations of the six critical neurons of the clique in the correct order and with the required delays to ignite a PB. The number of observed ABs was 66 % of the PBs, thus explaining why the average firing period of *eh*_3_ and *eh*_4_ was 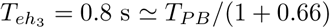, since their firing always triggered a PB or an AB.

To understand why in the case of ABs the sequential activation of the neurons of the clique did not lead to a PB ignition, we examined the value of synaptic resources for regular and aborted bursts, as shown in Fig. 4 (a). From the figure it is clear that 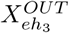 and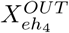 should reach a sufficient high value in order to observe a PB, otherwise one had an AB. Furthermore, the value reached by 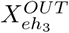 and 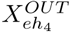 was related to the time passed from the last collective event and thus the requirement of a minimal value of the synaptic resources to observe a PB set a minimal value for the IGI, i.e. the interval between two PBs. As a matter of fact, as shown in Fig. 4 (b) the IGI values grew almost linearly with the values reached by 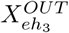 just before the PB, at least for 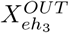 < 0.9. At larger values the relationship was no more linear and a saturation was observable, due to the fact that 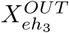 could not overcome one.

**Figure 4.**
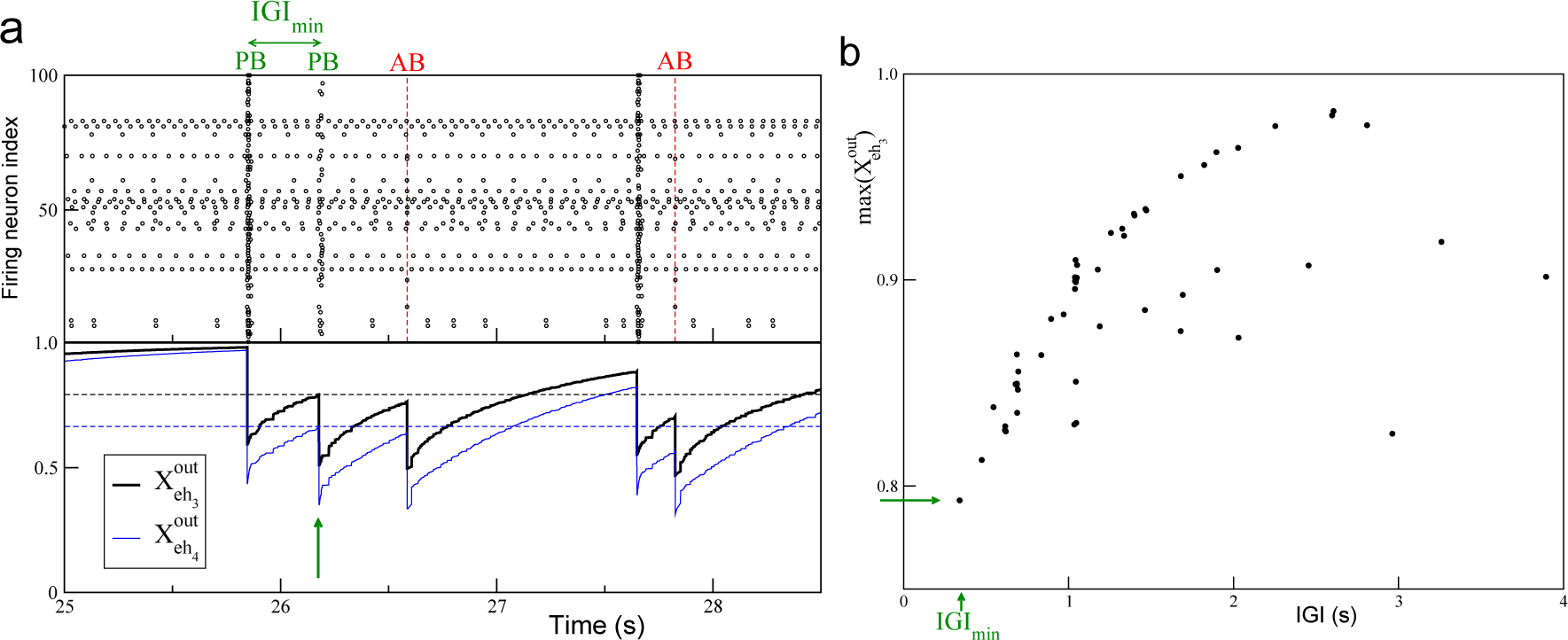
Model - Population bursts and synaptic resources. (a) Top panel: raster plot of the network activity, where population bursts (PBs) and aborted bursts (ABs) are shown. The vertical (red) dashed lines signal the occurrence of aborted burst. Bottom panel: average synaptic strength of the efferent connections of the two hub neurons *eh*_3_, *eh*_4_ in control conditions; the output effective synaptic strength is measured by the average value of the fraction 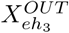 (thik black line), 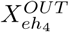 (thin blue line) of the synaptic transmitters in the recovered state associated to the efferent synapses. The dashed horizontal lines signals the values of the local maxima of 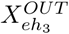 (black line, 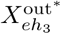 = 0.793) and 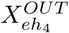 (blue line, 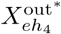 = 0.666) corresponding to the occurrence of the shortest IGI, IGI_min_. (b) Values of the local maxima of 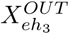 (max(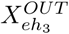)) in correspondence of the latest IGI. In both the figures the (green) arrow marks the occurrence of IGI_min_.

We could conclude that the slow firing of the couple (*eh*_3_, *eh*_4_), moderate by the inhibitory action of *ih*1 on *eh*3, was essential to ignite a PB, since a faster activity would not leave to the synapses the time to reach the minimal value required for a PB ignition, namely 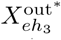 = 0.793 and 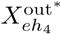 = 0.666. This could be better understood, by reconsidering the SND experiment on *ih*1, as expected the resection of neuron *ih*1 from the network led to a much higher activity of neurons *eh*3 and *eh*4, as shown in Fig. S6. However, this was not leading to the emission of any PBs, because in this case the value of 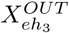 and 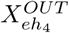 remained always well below the value required for a PB ignition.

### Tuning of PBs frequency upon hubs’ and LC cells’ stimulation

In order to better understand the role played by the hub and LC drivers for the collective dynamics of the network, we performed SNS experiments for a wide range of stimulation currents. The results of this analysis for currents in the range 14.5 mV ≤ *I*^stim^ ≤ 18 mV are shown in Fig. 5 (where all the driver hubs and six representative cases of driver LC cells are reported) and in Fig. 6 (a). The driver hub neurons could, upon SNS, usually lead to a reduction, or silencing, of PBs, apart for two cells (namely, *eh*_1_ and *eh*_4_) which, for specific stimulation currents, could even enhance the population activity. On the other hand, the 18 driver LC cells can be divided in two classes LC1 and LC2 according to their influence on the network dynamics upon SNS: a first group of 14 driver LC1 cells able mainly to reduce/stop the collective activity, and in few cases to increase the PB frequency, and a group of 4 LC2 neurons capable only to enhance the PB frequency. The three neurons *el*_1_, *el2* and *el*_3_, previously considered in subsection *Numerical evidences of driver LC cells* for comparison with experimentally identified LC cells, belonged to the class LC1 (see Fig. 1 and 2), while we have no experimental examples of LC2 cells.

**Figure 5.**
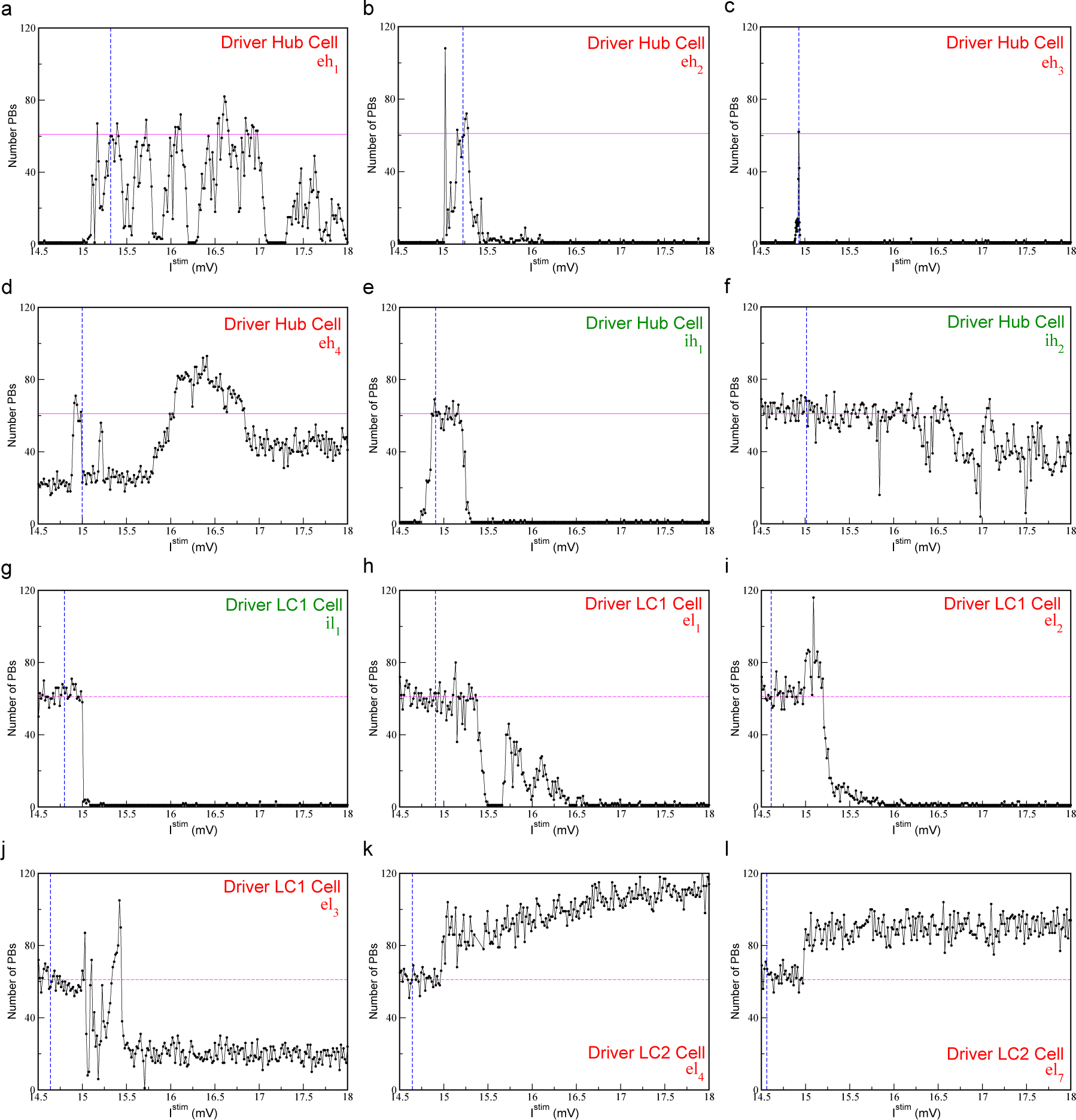
Model - PBs frequency is tuned by current stimulation of driver cells. The plots report the number of PBs emitted during SNS of the hub neurons *ih*_1_, *ih*_2_, *eh*_1,_ *eh*_2_, *eh*_3_, *eh*_4_ (a-f) as well for some driver LC cells (g-i) for a wide range of the stimulation current *I*^stim^ (over a time interval Δ*t* = 84 s). The blue vertical dashed lines, resp. the horizontal magenta solid line, refer to the value of the intrinsic excitability, resp. to the bursting activity, when the network is in control condition. The threshold value of the current is set to *V*_*Th*_ = 15 mV.

For what concerns the driver hubs’ dynamics, PBs were generated in the network whenever the hubs *eh*_2_ and *ih*_1_, both structurally connected to *eh*_3_, were stimulated with currents smaller than the excitability 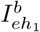 of the leader of the clique and within a specific interval (see Figs. 5 (b) and (e)). This means that in order to have a PB both neurons controlling *eh*_3_ should not fire faster than the leader of the clique. If this was not the case, the inhibition (originating from *ih*_1_) would not be anymore sufficient to balance the excitation (carried by *eh*_2_) or viceversa, thus leading *eh*_3_ to operate outside the narrow current window where it should be located to promote collective activity (see Fig. 5 (c)). In the case of *ih*_2_ and *eh*_4_ the SNS produced a less pronounced impact on the PB activity, their stimulation could never silence the network (as shown in Fig. 5 (d) and (f)), apart in two narrow stimulation windows for *ih*_2_. This is in agreement with what reported in Fig. 3 (a) for the SND, since the removal of neurons *eh*_4_ and *ih*_2_ only reduced the occurrence of PBs of 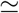 60%.

### LC drivers impact hub neurons

The SNS of LC1 drivers could induce, in 10 cases over 14 identified LC1 cells, a complete silencing in the network. A peculiar feature of eight out of these ten cases was that the PBs were completely suppressed as soon as these LC1 driver were brought supra-threshold: two examples are reported in Fig. 5 (g) and (i). The first example in Fig. 5 (g) refers to the unique inhibitory driver LC cell we have identified, namely *il*_1_, which was directly connected to the hub neuron *eh*_3_ (as shown in Fig. 3 (g)). A stimulation of *il*_1_ led to a decrease of the activity of *eh*_3_ and as a direct consequence of the PB activity. Fig. 5 (i) is devoted to *el*_2_, previously examined in Section *Numerical evidences of driver LC cells* and reported in Fig 1 (e.S-g.S). The depressive effect on the network activity due to the stimulation of *el*_2_, could be straightforwardly explained by the fact that *el*_2_ is directly connected to the inhibitory LC cell *il*_1_ and to the inhibitory hub driver *ih*_1_, thus performing an effective inhibitory action on the network, even if the stimulated driver *el*_2_ was excitatory. For the other six excitatory LC1 drivers acting on *il*_1_ only in two cases the PBs could not be completely blocked, and this happened when the cells were also directly connected to the hub driver *eh*_3_. In the two cases of LC1 driver cells able to block the population activity, but not acting through *il*_1_, these cells were exciting either *eh*_1_ or *ih*_1_, both belonging to the path with an inhibitory effect on *eh*_3_. The four remaining LC1 drivers that were able to reduce, but not to completely silence the population activity, acted either through *eh*_4_ (which was unable to block the PBs, even upon SND) or by simultaneously exciting and inhibiting *eh*_3_.

It should be remarked that nine of the previously discussed LC1 drivers could either enhance or reduce the PBs for different values of *I*^stim^. The double action of these neurons is exemplified by the two examples reported in Fig. 5 (h) and (j), which refer to neuron *el*_1_ and *el*_3_, already examined in connection with experimental data in Fig. 1 (b.S-d.S) and in Fig. 2 (a.S-f.S), respectively.

The neuron *el*_1_ was structurally connected to the inhibitory hub cell *ih*_1_, *el*_1_ was silent in control condition and once stimulated with a current *I*^stim^ = 15.135 mV was able to enhance population dynamics as shown in Fig. 1 (b.S-d.S). However, depending on the stimulation current it could even completely silence the PB activity, as shown in Fig. 5 (h). In order to understand in deeper details the mechanisms underlying both the enhancement and suppression of PBs, we stimulated *el*_1_ with *I*^stim^ ∈ [14.5 : 16] mV and for each value of the stimulation current we performed SND of the all neurons in the network. This analysis was aimed at identifying (for each value of *I*^stim^) the driver hub cells involved in the PB generation, i.e the neurons that upon SND reduced the population activity at least or more than 50%. The results of these experiments are shown in Fig. 6 (b-c), for sufficiently low stimulation currents (even above threshold) the activity of *el*_1_ had no influence at a network level, and this is consistent with the response of ih_1_ upon SNS reported in Fig. 5 (e). However for higher stimulation current the clique of functional hubs is modified by the action of *el*_1_: not all the hub cells previously identified remained relevant for the network activity and in some cases some new hub drivers was identified, as reported in Fig. 6 (b). The most significant modification is that the neuron *eh*_1_ was no more relevant (in most cases) for the PB generation, and this could be explained by the fact that the inhibitory hub ih1 is now controlled directly by *el*_1_. This is further confirmed by the fact that when the stimulation became sufficiently large the collective dynamics was completely silenced due to the high activity of *ih*_1_. As a matter of fact, some low activity in the network could be restored, due to a modified functional clique, for even larger current values above *I*^stim^ 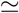 15.65 mV.

**Figure 6.**
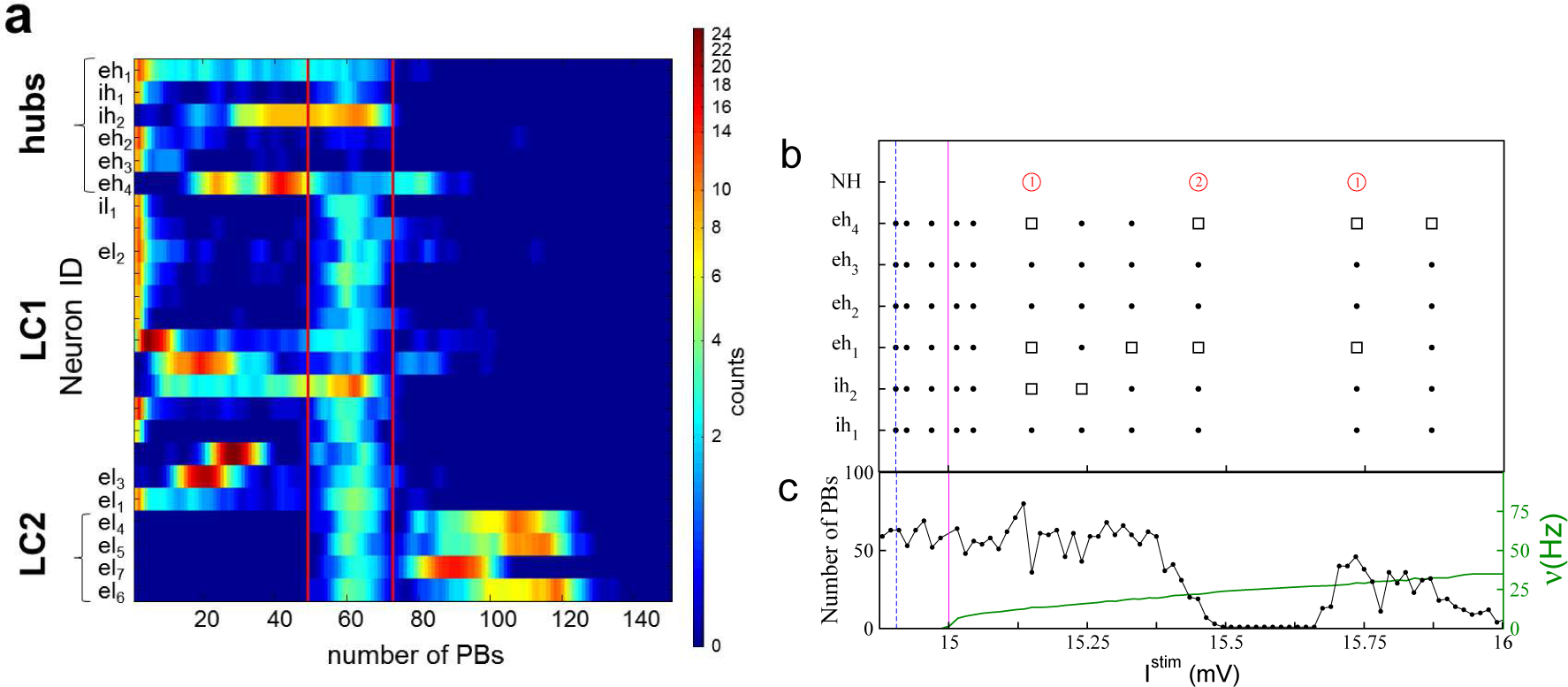
Model - Response to SNS of the driver cells. (a) Quantification of the response to SNS for each of the driver cells, sorted in three groups, respectivively hubs, LC1 and LC2. The heatmap displays how many times the SNS over a wide range of currents (namely, 14.5 mV ≤ *I*^stim^ ≤ 18 mV) induced a given number of PBs (x-axis) in the network. To facilitate the visualization, each row of the heatmap has been smoothed with a gaussian function of 1.58 standard deviation and unitary area. The red vertical lines denote the limits of activity in control condition: one standard deviation around the average. **Model - Current stimulation of driver LC cells can modify the functional clique of the network**. The panels (b),(c) refer to LC1 driver cell *el*_1_ (see Fig. 1 (b.S - d.S)). In the top panel (b) the configuration of the functional clique is reported for some sample stimulation current *I*^stim^ of *el*_1_. Full circles, resp. open squares, signal the presence, resp. absence, of the corresponding neuron of the functional clique *ih*_1_, *ih*_2_, *eh*_1_, *eh*_2_, *eh*_3_, *eh*_4_, while the open (red) circles indicate the presence of new neurons NH in the functional clique (the number of new neurons is reported inside the red circles). In the bottom panel (c) it is shown the number of PBs emitted by the network (black line with dots and left y-axis) and the firing frequency *v* of the LC cell (green line and right y-axis) during the current stimulation. The vertical (magenta) line marks the threshold value, *V*_th_, while the vertical (blue) dashed line signals the intrinsic excitability of the LC cell in control condition.

The LC1 driver *el*_3_ had also a double action leading to enhancement or depression of the collective activity as shown in Fig. 5 (j). This double action grounded in the following network architecture: *el*_3_ was structurally connected, via the bridge neuron *el*_1_, to *ih*_1_, whose impact on the network was to arrest the bursting apart a very narrow range of stimulation currents (Fig. 5 (e)); *el*_3_ was also structurally connected to *eh*_4_, which in some ranges of stimulation currents could enhance network dynamics (Fig. 5 (d)).

To conclude the analysis of the driver cells, we consider LC2 cells. These were excitatory neurons characterized by a low *I*^*b*^ (below the firing threshold *V*_th_) and, given the imposed correlation in the network model, by a high global structural connectivity (see Fig. S2). In control conditions these neurons were not active and did not participate to PBs. Two examples of the SNS of these neurons are reported in Figs. 5 (k) and (l) for LC2 drivers *el*_4_ and *el*_7_. Whenever they were stimulated above threshold they induced a sharp increase in the PB activity in the order of 50 %. Furthermore, in the case of LC2 drivers the SNS led in general to a more regular bursting dynamics, characterized by a smaller average Inter GDP Interval (of the order of the recovery time for the synapses, see *Methods* for details) and a smaller standard deviation (i.e. for *el*_7_ we measured < *IGI* >= 0.9 ± 0.6 s for *I*^stim^ = 15.9 mV) with respect to control conditions (where < *IGI* >= 1.4 ± 1.0 s). When LC2 cells were current stimulated, the action of enhancement of PBs activity was not mediated by the impact on other driver cells. As shown in Fig. S7, while the stimulation of LC1 drivers led to a noticeable modification of the firing rates of the hub cells, the SNS of LC2 driver cells had essentially no influence on the hubs. Therefore due to their high structural out-degree we can safely affirm that their influence on the network dynamics should be related to a cooperative excitatory effect. As a matter of fact, during the SNS of LC2 driver cells we observed the disappearence of ABs and as a consequence Inter GDP Interval become more regular, thus leading to an enhancement of the population activity. In particular, the disappearence of ABs was due to the fact that whenever eh3 was firing, a burst was emitted due to the presence of a higher level of excitation in the network, even when the synaptic resources of eh3 were below the minimal value required in control conditions, as discussed in the subsection *Synaptic resources and population bursts*.

## Discussion

We have developed a simple brain circuit model to mimic recent experimental results obtained in cortical slices of the mice Entorinhal Cortex during developmental stages [23]. These analysis revealed the existence of high and low functionally connected driver cells able to control the network dynamics. The fact that functional hubs can orchestrate the network dynamics is somehow expected [16,39], while the existence of driver neurons with low functional out-degrees has been revealed for the first time in [23]. In this paper, we focused mainly on the analysis of these latter class of drivers. which in control conditions were essentially irrelevant for the build-up of the GDPs. On the contrary, if single-handedly stimulated they could nevertheless strongly modify the frequency of occurrence of GDPs, as evident from the experimental findings reported in Figs. 1 (b-g.E) and 2 (a-f.E). In particular, their stimulation could lead both to an enhancement as well as to a reduction of the population activity (GDPs’ frequency). Quite remarkably, some of the driver LC cell were able to perform both these tasks as an effect of different stimulation frequencies as revealed by the experiment shown in Fig. 2 (a-f.E).

We have demonstrated that these experimental findings could be replicated in a simple spiking neural network model made of excitatory and inhibitory neurons with short-term synaptic plasticity and developmentally inspired correlations (see Figs. 1 (b-g.S) and 2 (a-f.S)). The analysis of the model has allowed to understand the fundamental mechanisms able to promote a single neuron to the role of network driver without being a functional hub, as usually expected.

In the model, all the driver neurons able to influence the network dynamics could be identified and they could be distinguished in neuronal hubs characterized by high out-degree or low functionally connected drivers. Functional hubs are highly excitable excitatory and inhibitory neurons arranged in a clique, whose sequential activation triggered the Population Bursts (analogous to GDPs). This in agreement with recent experimental evidences that small neuronal groups of highly active neurons can impact and control cortical dynamics [6–9]. On the other hand, driver LC cells are characterized by a lower level of excitability, but a higher structural connectivity with respect to driver hubs. Due to their low activity and functional connectivity in control conditions, these neurons were not fundamental for the PBs development, but were passively recruited during the burst, or even completely silent. The LC drivers can be divided in two classes LC1 and LC2 according to their influence on the population activity whenever stimulated with different values of DC current: the majority of them were able both to enhance and reduce (or even set to zero) the frequency of occurrence of the PBs (LC1), while a small group was able only to enhance the PBs’ frequency with respect to control conditions (LC2). Noticeably, driver LC1 cells were structurally connected to the hubs (directly or via a bridge LC cell). Therefore, whenever stimulated they can influence the network activity by acting on the clique dynamics. In most cases, even if these cells were excitatory, their action on the network was mainly depressive, since either they stimulated directly inhibitory hubs or the inhibitory LC1 driver, which acted as a bridge over the clique. In more than the 50% of the cases (8 over 14) whenever brought over threshold driver LC1 cells led to a complete arrest of the PB activity.

Driver LC2 cells instead were silent in control conditions and highly structurally connected, therefore they were putative structural hubs. As a matter of fact, whenever brought supra threshold they favoured a more regular collective dynamics. The activation of the many efferent connections of LC2 drivers led to the creation of many alternative pathways for the PB igniton, in a sort of homeostatic regulation of the network which led to an optimal employ of the synaptic resources [40] with the corresponding disappearence of the aborted bursts, largely present in control conditions.

Furthermore, we have shown that the stimulation of single driver LC cells was not only able to alter the collective activity but also to deeply modify the role of neurons in the network, such that some neurons can be promoted to the role of driver functional hubs or driver hubs can even loose their role (see Fig. 6 (b-c)). At variance with purely excitatory networks [11], the synchronized dynamics of the present network, composed of excitatory and inhibitory neurons, is less vulnerable to targeted attacks to the hubs [41,42]. As demonstrated by the fact that different firing sequences of hub neurons can lead to population burst ignitions (see Fig. 3 (e)) and that hubs can be easily substituted in their role by LC driver cells when properly stimulated.

Another relevant aspect is that the inclusion of inhibitory neurons in the network did not cause a trivial depressing action on the bursting activity, as it could be naively expected, but instead they can play an active role in the PB build-up. Our analysis clearly demonstrate that their presence among the driver cells is crucial in determining and controlling the PB activity, somehow similarly to what found in [19] where it has been shown that the emergence of sharp-wave in adult hippocampal slices was controlled by single perisomatic-targeting interneurons.

Our results suggest that inhibitory neurons can have a major role in information encoding by rendering on one side the population dynamics more robust to perturbations of input stimuli and on another side much richer in terms of possible repertoire of neuronal firings. These indications confirm the key role of inhibitory neurons in neural dynamics, already demonstrated for the generation of brain rhytms [43,44] and for attentional modulation [45].

Recently there has been a renovated interest on the existence and role of neuronal cliques within the brain circuitries [12,46]. Cliques have been proposed as structural functional multiscale computational units whose hierarchical organization can lead to increasingly complex and specialized brain functions and can ground memory formations [46]. In addition, activation of neuronal cliques as in response to external stimuli or feedforward excitation can lead to a cascade of neuronal network synchronizations with distinct spatio-temporal profiles [12]. Our results provide a further understanding on how cliques can emerge (spontaneously during development) and modify (in response to stimuli similarly to the SNS here discussed) with a consequent reshaping of the spatio-temporal profile of the dynamics of the network in which the clique is embedded. Notably, it is the presence of inhibitory neurons within the network to favour the emergence of different cliques by empowering drivers with different functional connectivity degree. While functional driver hubs guarantee the functioning of the network synchronization in absence of stimuli (such as during development and in non-stimulated conditions), LC drivers widen the ability of the network to play distinct synchronization profiles (i.e. spatio-temporal activations) possibly underlying emergent functions within the brain networks.

Finally, our results could be of some relevance also for the control of collective dynamics in complex networks [47,48]. Usually the controllability of complex networks is addressed with linear dynamics [49,50]. However, at present there is not a general framework to address controllability in nonlinear systems, in this context the SND and SNS protocols we developed for pulse-coupled networks could be extended to general complex networks as a tool to classify driver nodes and as a measure of controllability [51].

## Materials & Methods

### Experiment

#### Animal Treatment

All animal use protocols were performed under the guidelines of the French National Ethic Committee for Sciences and Health report on “Ethical Principles for Animal Experimentation” in agreement with the European Community Directive 86/609/EEC. Double-homozygous Mash1BACCreER/CreER/RCE:LoxP+/+ and Dlx1/2CreER/CreER/ RCE:LoxP+/+ [52,53] male mice were crossed with 7 to 8-week-old wild-type Swiss females (C.E Janvier, France) for offspring production. To induce CreER activity,we administered a tamoxifen solution (Sigma, St. Louis, MO) by gavaging (force-feeding) pregnant mice with a silicon-protected needle (Fine Science Tools, Foster City, CA).

#### Slice preparation and calcium imaging

Horizontal cortical slices (400 mm thick) were prepared from 8 day old (P8) GAD67^gfp^ (n = 29), Lhx6^iCre^::RCE:LoxP (n = 23) or 5-HT3aR-BAC^EGFP^ (n=15) mouse pups with a Leica VT1200 S vibratome using the Vibrocheck module in ice-cold oxygenated modified artificial cerebrospinal fluid (0.5 mM CaCl_2_ and 7 mM MgSO_4_; NaCl replaced by an equimolar concentration of choline). Slices were then transferred for rest (1 hr) in oxygenated normal ACSF containing (in mM): 126 NaCl, 3.5 KCl, 1.2 NaH_2_PO_4_, 26 NaHCO_3_, 1.3 MgCl_2_, 2.0 CaCl_2_, and 10 D-glucose, pH 7.4. For AM-loading, slices were incubated in a small vial containing 2.5 ml of oxygenated ACSF with 25 ml of a 1 mM Fura2-AM solution (in 100% DMSO) for 20-30 min. Slices were incubated in the dark, and the incubation solution was maintained at 35-37C^0^. Slices were perfused with continuously aerated (3 ml/min; O_2_/CO_2_-95/5%) normal ACSF at 35-37 C^0^. Imaging was performed with a multibeam multiphoton pulsed laser scanning system (LaVision Biotech) coupled to a microscope as previously described (see [54]). Images were acquired through a CCD camera, which typically resulted in a time resolution of 50-150 ms per frame. Slices were imaged using a 203, NA 0.95 objective (Olympus). Imaging depth was on average 80 mm below the surface (range: 50-100 mm).

#### Experimental Design

A total of n=67 neurons were electrophysiologically stimulated and recorded following the criteria: (1) stable electrophysiological recordings at resting membrane potential (i.e., the holding current did not change by more than 15 pA); (2) stable network dynamics measured with calcium imaging (i.e., the coefficient of variation of the inter-GDP interval did not exceed 1); (3) complete labeling of the recorded cell; and (4) good quality calcium imaging while recording. Neurons were held in current-clamp using a patch-clamp amplifier (HEKA, EPC10) in the whole-cell configuration. Intracellular solution composition was (in mM): 130 K-methylSO_4_, 5 KCl, 5 NaCl, 10 HEPES, 2.5 Mg-ATP, 0.3 GTP, and 0.5% neurobiotin. No correction for liquid junction potential was applied. The osmolarity was 265-275 mOsm, pH 7.3. Microelectrodes resistance was 6-8 MOhms. Uncompensated access resistance was monitored throughout the recordings. Recordings were digitized online (10 kHz) with an interface card to a personal computer and acquired using Axoscope 7.0 software. Spontaneous EPSPs were detected and analyzed using the MiniAnalysis software. For most stimulation experiments, the movie acquisition time was separated evenly in three epochs: (1) a 2 min resting period during which the cell was held close to ***V_res_t*** (i.e., zero current injection); (2) a 2 min stimulation period during which phasic stimulation protocols were applied; and (3) a 2 min recovery period where the cell was brought back to resting membrane potential. Stimulation protocol: suprathreshold current pulses (amplitude: 100-200 pA, duration: 200 ms) repeated at one of the following frequencies: *v*_*S*_ =*IGI*/3 or IGI/2, where IGI is the average frequency of GDP occurrence under control conditions.

#### Analysis of the Data

**Signal Detection** We used custom designed MATLAB software [55] that allowed: (1) automatic identification of loaded cells; (2) measuring the average fluorescence transients from each cell as a function of time; (3) detecting the onsets and offsets of calcium signals, and (4) reconstructing the functional connectivity of the imaged network.

**Statistical Analysis** Network synchronizations (GDPs) were detected as synchronous onsets peaks including more neurons than expected by chance, and their time stamp denoted by *t*_G_. The Inter-GDP-interval (IGI) is defined as the interval between two consecutive GDPs. To establish whether the stimulation of a single neuron was able to influence the frequency of GDPs occurrence, we first calculte the average IGI in the three epochs: pre-stimulus (control), during the stimulation period, and poststimulus. Due to the variability distribution of IGI in each interval, we calculate, the average IGI in a window of *t*_s_ = 60s calculated starting from each *t*_G_, eliminating the data corresponding to overlaps between epochs. To test for the significance of the change in the period of GDP due to single neuron stimulation a Kolmogorov-Smirnov test is applied between all the 3 resulting distributions of average IGI and a significance level of *p* < 0.05 is chosen to be realiable.

Also, for the *i*-th GDP, a phase measure Φ*i* is defined respect to the control IGI as follows:

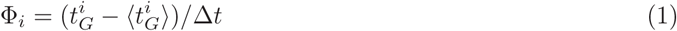

where Δ*t* is the average IGI interval in the control condition, and 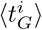 = *i* × Δ*t* is the expected occurrence if the *i*-th GDP, according to the control condition.

### Model

#### Definition of the model

To study the response of bursting neural networks to single neuron stimulation and removal, we employed the Tsodyks-Uziel-Markram (TUM) model [32]. Despite being sufficiently simple to allow for extensive numerical simulations and theoretical analysis, this model has been fruitfully utilized in neuroscience to interpret several phenomena [33,34,56]. We have considered such a model to mimic the dynamics of developing brain circuitries, which is characterized by coherent bursting activities, such as *giant depolarizing potentials* [16,57]. These coherent oscillations emerge, instead of abnormal synchronization, despite the fact that the GABA transmitter has essentially an excitatory effect on immature neurons [28].

In this paper we consider a network of N leaky-integrate-and-fire (LIF) neurons interacting via synaptic currents regulated by short-term synaptic plasticity (depression and facilitation) according to the model introduced in [32]. In particular the facilitation mechanism is present only for synapses targeting inhibitory neurons.

The time evolution of the membrane potential *Vi* of each neuron reads as

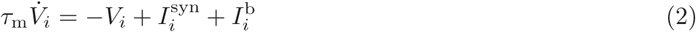

where *τ*_m_ is the membrane time constant, 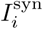 is the synaptic current received by neuron *i* from all its presynaptic inputs and 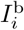 represents its level of intrinsic excitability. The membrane input resistance is incorporated into the currents, which therefore are measured in voltage units (mV).

Whenever the membrane potential 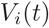 reaches the threshold value *V*_th_, it is reset to *V*_r_, and a spike is sent towards the postsynaptic neurons. For the sake of simplicity the spike is assumed to be a *δ*-like function of time. Accordingly, the spike-train *S*_*j*_ (t) produced by neuron *j*, is defined as,

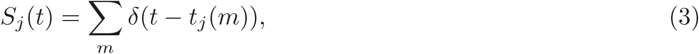

where *t*_*j*_ (*m*) represent the *m*-th spike time emission of neuron *j*. The transmission of the spike train *S*_*j*_ to the efferent neurons is mediated by the synaptic evolution. In particular, by following [58] the state of the synaptic connection between the jth presynaptic neuron and the ith postsynaptic neuron is described by three adimensional variables, *X*_*ij*_, *Y*_*ij*_, and *Z*_*ij*_, which represent the fractions of synaptic transmitters in the recovered, active, and inactive state, respectively and which are linked by the constraint *X*_*ij*_, +*Y*_*ij*_, + *Z*_*ij*_ = 1. The evolution equations for these variables read as

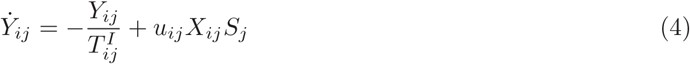

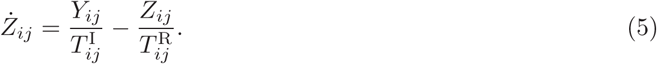

Only the active transmitters react to the incoming spikes *S*_*j*_ : the adimensional parameters *u*_*ij*_ tune their effectiveness. For the synapses targeting excitatory neurons *u*_*ij*_ ≡*U*_*ij*_ stay constant, while for the synapses targeting inhibitory neurons *u*_*ij*_ display a dynamical evolution (facilitation)

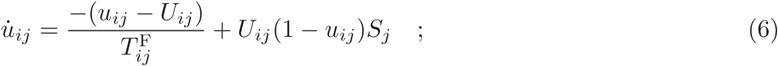

where 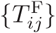 control the decay of the facilitation variables. Moreover, 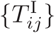 represent the characteristic decay times of the postsynaptic current, while 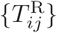 are the recovery times from synaptic depression.

Finally, the synaptic current is expressed as the sum of all the active transmitters (post-synaptic currents) delivered to neuron *i*

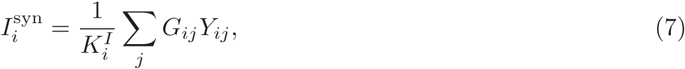

where *G*_*ij*_ are the coupling strengths, whose values can be finite (zero) if the presynaptic neuron *j* is connected to (disconnected from) the postsynaptic neuron *i*. Furthermore, if the presynaptic neuron is excitatory (inhibitory) the sign of *G*_*ij*_ will be positive (negative).

In this paper, we consider a *diluted* network made of N = N_e_ + *N*_*i*_ = 100 neurons, where *N*_e_ = 90 (*N*_*i*_ = 10) is the number of excitatory (inhibitory) cells. The *i*-th neuron has 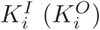 afferent (efferent) synaptic connections distributed as in a directed Erdös-Rényi graph with average in-degree 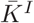 = 10, as a matter of fact also the average out-degree was 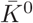 = 10. The sum appearing in (7) is normalized by the input degree 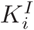 to ensure homeostatic synaptic inputs [59,60].

The propensity of neuron *i* to transmit (receive) a spike can be measured in terms of the average value of the fraction of the synaptic transmitters 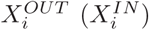 in the recovered state associated to its efferent (afferent) synapses, namely

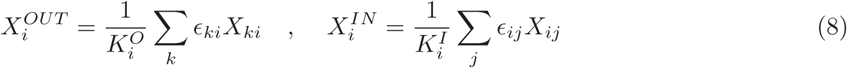

where ϵ_*ij*_ is the connectivity matrix whose entries are set equal to 1 (0) if the presynaptic neuron j is connected to (disconnected from) the postsynaptic neuron i.

The intrinsic excitabilities of the single neurons 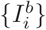 are randomly chosen from a flat distribution of width 0.45 mV centered around the value *V*_th_ = 15 mV, with the constraint that 10% of neurons are above threshold. This requirement was needed to obtain bursting behavior in the network. With this choice the distribution of the single neuron firing rates under control conditions is in the range [0.05; 22] Hz.

Furthermore, we have considered networks where a negative correlation between the intrinsic neuronal excitability 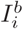 and the total connectivity (in-degree plus out-degree) 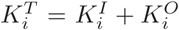 is embedded. To generate this kind of correlation the intrisic excitabilities are randomly generated, as explained above, and then assigned to the various neurons accordingly to their total connectivities 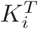, thus to ensure an inverse correlation between 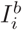 and 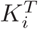. The correlation is visualized in Fig S2.

For the other parameters, we use the following set of values: *τ_m_* = 30 ms, *V*_r_ = 13.5 mV, *V*_th_ = 15 mV. The synaptic parameters 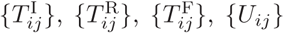 and 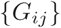 are Gaussian distributed with averages 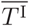 = 3 ms, 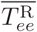 = 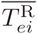 = 800 ms, 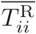 = 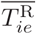 = 100 ms 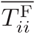 = 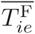 = 1000 ms, 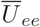 = 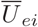 = 0.5, 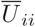 = 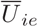 = 0.04, and 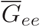 = 45 mV, 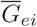 _*ei*_= 135 mV, 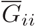 = 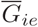 = 180 mV respectively, and with standard deviation equal to the half of the average. These parameter values are analogous to the ones employed in [32] and have a phenomenological origin.

In order to have an accurate and fast integration scheme, we transformed the set of ordinary differential equations (2), (4), (5) and (6) into an event-driven map [61] ruling the evolution of the network from a spike emission to the next one (see Supplementary Note 1 for more details on the implementation of the event-driven map). It is worth to stress that the event-driven formulation is an exact rewriting of the dynamical evolution and that it does not involve any approximation.

#### Population Bursts and Aborted Bursts

In order to identify a population burst we have binned the spiking activity of the network in time windows of 10 ms. A population burst is identified whenever the spike count involves more than 25 % of the neural population. In order to study the PB build up, a higher temporal resolution was needed and the spiking activity was binned in time windows of 1 ms. The peak of the activation was used as time origin (or center of the PB) and it was characterized by more than 5% of the neurons firing within a 1 ms bin. The time window of 70 ms preceding the peak of the PB was considered as the build up period for the burst. In particular, the threshold crossing times have been defined via a simple linear interpolation based on the spike counts measured in successive time bins.

These PB definitions gave consistent results for all the studied properties of the network. The employed burst detection procedure did not depend significantly on the precise choice of the threshold, since during the inter-burst periods only 17 - 20 % of neurons were typically firing, while 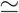80 % of the neuronal population contributed to the bursting event.

The average interburst interval for the network with (without) correlations under control conditions was 1.4 ± 1s (0.3 ± 0.1 s), while the burst duration was 17 ± 3 ms (30 ± 6 ms) for a network made of N =100 neurons.

Aborted bursts were collective events associated to an observable enhancement of the network activity, but well below the threshold we fixed to detect PBs. The period of occurrence of the ABs was comparable to the average interburst interval, while their number corresponded to the 66 % of the PBs in the network with correlations and to the 47 % in random networks.

#### Functional Connectivity

In order to highlight statistically significant time-lagged activations of neurons, for every possible neuronal pair we measured the cross-correlation between their spike time series. On the basis of this crosscorrelation we eventually assign a directed functional connection among the two considered neurons, similarly to what reported in [16,62] for calcium imaging studies.

Let us explain how we proceeded in more details. For every neuron, the action potentials timestamps were first converted into a binary time series with one millisecond time resolution, where ones (zeros) marked the occurrence (absence) of the action potentials. Given the binary time series of two neurons *a* and *b*, the cross correlation was then calculated as follows:

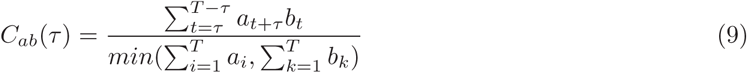

where {*a*_*t*_},{*b*_*t*_} represented the considered time series and *T* was their total duration. Whenever *C*_*ab*_(*τ*) presented a maximum at some finite time value *τ _max_* a functional connection was assigned between the two neurons: for *τ _max_* < 0 (*τ* _max_ > 0) directed from a to b (from b to a). A directed functional connection cannot be defined for an uniform cross-correlation corresponding to uncorrelated neurons or for synchronous firing of the two neurons associated to a Gaussian C_*ab*_(*τ*) centered at zero. To exclude the possibility that the cross correlation could be described by a Gaussian with zero mean or by a uniform distribution we employed both the Student’s t-test and the Kolmogorov-Smirnov test with a level of confidence of 5%. The functional out-degree 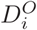 (in-degree 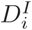) of a neuron *i* corresponded to the number of neurons which were reliably activated after (before) its firing.

*Time series surrogates*

In order to treat as an unique event multiple spike emissions occurring within a PB, different time series surrogates were defined for different kind of analysis according to the following procedures:

1. for the definition of the functional in-degree 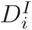 and out-degree 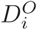, all the spiking events associated to an inter-spike interval longer than 35 ms were considered. Since we observed that this was the minimal duration of an inter-spike outside a PB and it was larger than the average duration of the PBs. This implies that for each neuron only the timestamp of the first spike within a PB was kept;
2. for the description of the PBs build up only the timestamps of the first action potential emitted within a window of 70 ms preceding the PB peak was taken into account;
3. for the analysis of the network activity during inter-burst periods, all action potentials emitted out of the PBs were considered.

## Acknowledgments

The authors had useful interactions with M. Di Bernardo, S. Olmi, M. Timme. This work has been realized within the activity of the Joint Italian-Israeli Laboratory on Integrative Network Neuroscience financed by the Italian Ministry of Foreign Affairs (S.L., P.B., A.T.). The authors acknowledge financial support from the European Commission under the program “Marie Curie Network for Initial Training”, project N. 289146, “Neural Engineering Transformative Technologies (NETT)” (S.L.,D.A.-G., A.T.), by the A* MIDEX grant (No. ANR-11-IDEX-0001-02) and by the I-Site Paris Seine Excellence Initiative (No. ANR-16-IDEX-0008), both funded by the French Government programme “Investissements d’Avenir” (A.T. and D.A.-G.), from Ikerbasque (The Basque Foundation for Science) (P.B.) and from the Ministerio Economia, Industria y Competitividad of Spain (grant SAF2015-69484-R) (P.B.). The authors declare no competing financial interests.

## Supporting Information

**Figure S1:**

**Experimental Setup**. (a) Slice of Enthorinal cortex with Calcium indicator. Contoured cells are the active cells. (b) Single neuron activity as a function of time during the three phases of the stimulation protocol: 1) pre-stimulation period where only spontaneous activity is recorded (2 min.); 2) single neuron is injected with pulses of fixed amplitude at a certain frequency *ν s* (2 min.); 3) post stimulus period without stimulation (2 min.). (c) Calcium trace for a selected neuron during the whole protocol. A time point is plotted in the upper part of the calcium trace whenever an onset of activity is present. Red (blue) traces denotes stimulation (control) epochs.

**Figure S2:**

**Model - Setup for connectivity and excitability**. (a) Negative correlation between intrinsic excitability *I*^*b*^ and total connectivity *K*^T^. The (magenta) line indicates the threshold value *V*_*th*_ = 15 mV, dividing supra-threshold from sub-threshold neurons. (b) Scatter plot of the in-degrees and out-degrees for each neuron in the network (no correlation). In both the figures dots (asteriskes) refer to excitatory (inhibitory) neurons. The data refer to *N* = 100 and all the parameter values are defined as in Methods.

**Figure S3:**

**Model - Response of the newtork without correlations to single neuron deletion (SND) and stimulation (SNS).** (a), (b) Number of PBs recorded during SND (SNS) experiments versus the label of the removed (stimulated) neurons, ordered accordingly to their average firing rates *ν* under control conditions (shown in panel c)). During SNS experiments each neuron was stimulated with a DC step *I*^stim^ = 15.90 mV for a time interval Δ*t* = 84 s. The horizontal dashed line shows the average number of PBs emitted in control conditions within an interval Δ*t* = 84 s, while the horizontal dotted lines mark the 50% variation. The vertical dashed red line separates firing neurons (on the right side) from silent neurons (on the left side) in control conditions. In all the panels, dots (asteriskes) symbols refer to excitatory (inhibitory) neurons.

**Figure S4:**

**Model - Structural properties of the neurons.** Scatter plots showing the structural properties of the neurons of the network in control conditions, (a) intrinsic excitability *I*^b^, (b) total structural connectivity *K*^T^. Dots (asteriskes) symbols refer to excitatory (inhibitory) neurons. The critical neurons *ih*_1_, *ih*_2_, *eh*_1_, *eh*_2_, *eh*_3_, *eh*_4_, belonging to the functional clique responsible for the PB-build up, are signaled by open circles, while the driver LC cells are denoted by open squares, in both cases red (green) contour codes for excitatory (inhibitory) neurons. The vertical dashed line separates firing neurons (on the right side) from silent neurons (on the left side) in control conditions, while the horizontal (magenta) line marks the threshold value, *V*_*s*_ = 15 mV, dividing supra-threshold from sub-threshold neurons. The neurons are ordered accordingly to their average firing rate in control conditions.

**Figure S5:**

Model - The activity of driver hub cells. Cross correlation functions between the hub drivers. The blue histograms are calculated using the first spike fired by each neuron during the PBs build-up. The red histograms are calculated using the spikes fired out of the PBs and the ABs. Note that during the PB build-up, neurons activate reliably in the following order *eh*_1_ → *ih*_1_ → *ih*_2_ → *eh*_2_ → *eh*_3_→*eh*_4_. During the out-of-burst activity, identical time lagged activation are preserved among the structurally connected pairs, namely *eh*_1_ → *ih*_1_ → *ih*_2_ → *eh*_2_ and *eh*_2_ → (*eh*_3_, *eh*_4_).

**Figure S6:**

**Model - Deletion (SND) of the inhibitory neuron** *ih*_1_ **of the clique leads to the arrest of the bursting activity**. In the top panel raster plot of the network activity during SND experiment on ih_i_ (neurons are labeled accordingly to the natural index); dots (asteriskes) symbols refer to excitatory (inhibitory) neurons, while large dots and dashed (red) lines refer to the driver hubs *eh_3_, *eh*_4_*. Bottom panel: average synaptic strength of the efferent connections of the two driver hubs neurons *eh_3_, *eh*_4_;* the output effective synaptic strength is measured by the average value of the fraction 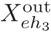 (black line), 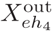 (blue line) of the synaptic transmitters in the recovered state associated to the efferent synapses. The output effective synaptic strengths are always under the minimal values for PB ignition 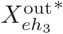, 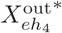 represented by the dashed lines (see also Fig.4 in the main text).

**Figure S7:**

**Model - Results of the SNS of LC drivers on the firing activity of the neurons of the clique**. Firing frequency *ν* of the neurons of the clique versus the current stimulation *I*^stim^ during SNS of LC drivers *el*_1_ (a) and *el*_7_ (b): (brown) stars, (red) crosses, (maroon) squares, (black) points, (green) diamonds, (blue) triangles refer respectively to *eh*_1_, *eh*_2_, *eh_3_, *eh*_4_, ih*_1_, *ih*_2_. The vertical (magenta) line marks the threshold value Vt_h_, while the (black) arrows signal the firing frequency of the neurons of the clique in control condition 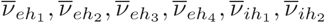

**Table S1:**

**Model - Routes leading to PBs.** Spike time delays *ΔΤ* between two successive firing of the neurons forming the functional clique along the main and secondary route leading to bursting. Neurons *eh*_3_ and *eh*_4_ are assumed to fire essentially at the same time, since *eh*_4_ fires almost immediately after *eh*_3_ within 0.03 - 0.04 ms. Notice that *eh*_1_ is the only hub driver cell firing twice before a PB: the two routes could be distinguished by the time occurrence of the second spike of *eh*_1_ and this event is denoted by an asterisk in the table. The arrows indicate the order of firing in the sequence.

